# A combined computational and experimental approach to successfully predict the behavior of metastatic tumor-associated macrophages in human breast cancer

**DOI:** 10.1101/2025.02.06.636799

**Authors:** Amel Bekkar, Angélique Pabois, Isaac Crespo, Riccardo Turrini, Ioannis Xenarios, Marie-Agnès Doucey

## Abstract

We identified in human breast cancer a scarce population of Tumor-Associated Monocytes (TAMs) endowed with a pro-metastatic activity and associated with reduced distant-metastasis free survival of patients. We developed a novel framework combining computational and experimental methodologies to dampen TAM pro-metastatic activity. Multi-modal experimental data from TAMs exposed in vitro to a series of perturbations were collected to build a Boolean dynamical network of TAMs. This framework successfully identified the biological pathways underlying TAM pro-metastatic activity and predicted potent pharmacological interventions that inhibited the pro-metastatic activity of TAMs isolated from patient tumors. This study showcases the power of integrating computational predictions with experimental validation in identifying new therapeutic avenues that can be extended to other cancer types.

## Introduction

Metastasis is the multiple process by which an original primary tumor develops into a distal secondary tumor. It remains the most life-threatening risk factor for cancer patients and the leading cause of mortality in breast cancer with 20-25% of European patients developing distant metastases post-treatment and eventually succumbing to the disease ^1,2^. In breast cancer and many other cancer types, clinical evidence demonstrates that lymph node metastasis is a critical prognostic indicator for systemic spread and reduced survival^3,4^. Pro-metastatic TAMs promote breast cancer metastasis to lymph nodes and distant sites by protecting them from destruction by the immune system, contributing to resistance to therapy, and inducing Epithelial-Mesenchymal Transition (EMT) i.e. tumor cell acquisition of more aggressive characteristics^2,5^. However, the underlying cellular and molecular mechanisms are not yet fully elucidated. Understanding the nature and mechanism by which TAMs induce breast cancer metastasis will facilitate the early diagnosis, prognosis, and treatment of patients with metastatic breast cancer^2,5^.

We have identified in human breast cancer a specific population of monocytes that represents less than 5% of total TAMs and is endowed with a strong pro-metastatic activity. Surprisingly, these TAMs were found exclusively inserted into tumor lymphatics^6^ and associated with lymph node metastasis in breast cancer patients. Further, we have identified that these pro-metastatic TAMs specifically express a cell surface protein required for their insertion into lymphatics and that functions as a marker of this TAMs population (undisclosed marker and thereafter called pro-metastatic marker).

To understand the molecular bases underlying the pro-metastatic activity of this scarce population of TAMs, we constructed and validated a dynamical computational model of these cells based on experimental multi-modal data^7,8^. Experimental assessment of the model predictions revealed that the model was capable of reliably simulating and predicting TAM pro-metastatic activity. Further, the model simulations identified a complex interplay of critical biological pathways underlying TAM pro-metastatic activity that would have been impossible to address experimentally. Predicting potent combinations of therapies to impair tumor growth and metastasis is a major challenge in the field of systems biology given the number of biological pathways to be investigated in the absence of prior knowledge and the complexity and heterogeneity of patient tumor ecosystems.

While combination therapy emerges as a powerful approach to enhance cancer patient clinical benefit, the daunting magnitude of combinations to be explored precludes their experimental and clinical testing. Hence, leveraging computational models to predict potent combinations of therapies for cancer patients represents an opportunity to address the unmet need for a rational approach to combination therapy; at the same time, it is a major challenge given the limited methodologies developed to date to assemble relevant experimental and computational approaches. To investigate the molecular interactions driving TAM-induced metastasis, we employed a logic-based modeling approach, which leverages Boolean variables to represent molecular states and interactions. Logic models have become increasingly valuable in oncology and systems biology for simulating complex biological networks ^9–12^, especially when detailed kinetic parameters are not available, as they allow researchers to capture essential signaling dynamics with minimal data requirements. This approach facilitates an integrative experimental and computational framework of hypothesis generation and validation. Furthermore, logic modeling has proven particularly adept at identifying novel therapeutic targets and combination treatments by highlighting pathway interdependencies. However, evidence of the contribution of logic models to pharmacology is yet lacking. Indeed, such models have not been validated experimentally, particularly within complex ecosystems such as patient tumors. Here we report, to the best of our knowledge, the first successful attempt to identify highly potent combinations of treatments that dampened patient TAM pro-metastatic activity. The successful outcome of these methodologies paves the way for a rational approach to combination therapy in oncology.

## RESULTS

### Biological approach to the understanding of patient TAM pro-metastatic activity

In human breast cancer, TAMs represent up to 60% of the immune infiltrate with, however, a large variability. We show that in breast tumors and patient peripheral blood, monocytes endowed with a pro-metastatic activity expressed at their surface a specific marker (pro-metastatic marker, manuscript in preparation) which allows their insertion into tumor lymphatics^6^. TAMs expressing this marker are found in all patients with invasion of tumor cells to the lymph node at the time of diagnosis and in 50% of the patients without detectable lymph node metastasis (manuscript in preparation). Hence, pro-metastatic TAMs represent a low fraction of breast tumor TAM (about 5%), are found inserted into tumor lymphatics, and are capable of promoting tumor cell spreading by inducing the transmigration of tumor cells through the lymphatic endothelium. They are likely to function as a gateway for tumor cell spreading and detach from the lymphatic vessel walls to reach the peripheral blood where they are detectable at very low frequencies. Importantly, we report that exposure in vitro of circulating monocytes from healthy donors to a conditioned medium of metastatic tumor cells induced the expression of the pro-metastatic marker in 20% of monocytes. These in vitro-derived pro-metastatic monocytes share a comparable functional phenotype and gene expression profile with pro-metastatic patient TAM and represent an excellent model system (manuscript in preparation) that we have used in this study. We established a tumor cell transmigration assay which serves as a robust functional metric of TAM pro-metastatic activity (manuscript in preparation). This assay mimics the process of tumor cell transmigration through the lymphatic endothelium mediated by pro-metastatic TAMs (Figure 1, scheme).

**Figure 1.**
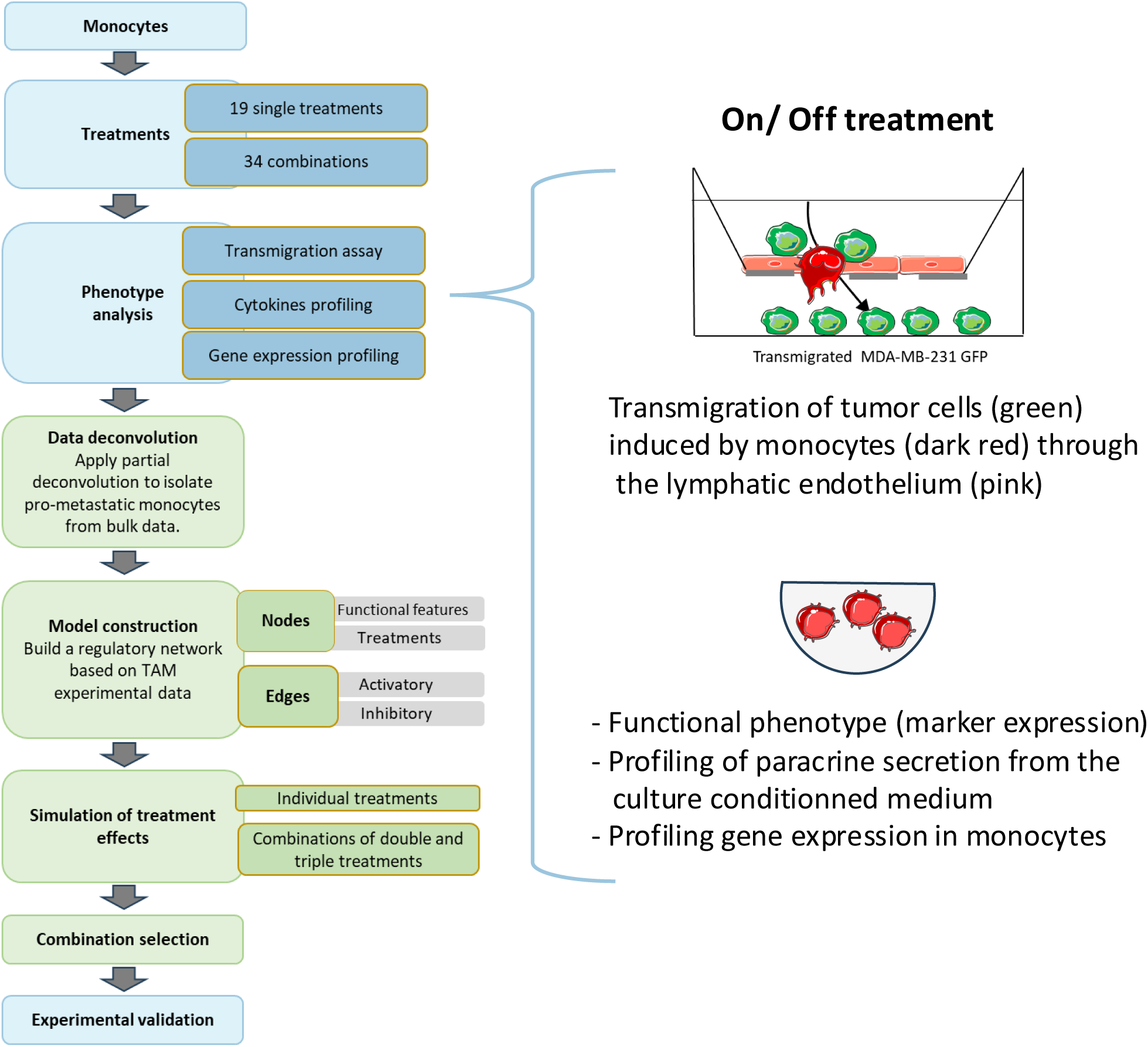
Workflow diagram of assembled experimental and computational approaches to simulate and predict TAM pro-metastatic activity. Experimental and computational approaches are depicted in blue and white, respectively. TAM pro-metastatic activity was measured with the tumor cell transmigration assay depicted on the right end.

### Successful assembly of experimental and computational modeling approaches to generate a dynamical network of pro-metastatic TAMs

We implemented a multi-step experimental and computational approach to evaluate the effects of various treatments on monocyte populations and their influence on tumor cell transmigration (Figure 1). Monocytes were treated with 19 individual treatments and 34 combinations, followed by comprehensive phenotypic analysis, including transmigration assays, cytokine profiling, and gene expression profiling (Table 1). We inferred the dynamical network of the pro-metastatic monocytic population from bulk monocyte experimental data (Figure 2) after partial data deconvolution to isolate pro-metastatic monocytes from bulk data, enabling more precise profiling of their functional phenotype. The attractor states were evaluated under all single, dual, and triple treatment conditions. By integrating in silico predictions of node states with their corresponding Orthogonal Partial Least Squares (OPLS) weights (Figure 1 and 3A), we computed a transmigration score to quantify pro-metastatic activity (Method section). This score allowed us to rank treatments and their combinations relative to Tumor Conditioned Medium (TCM), which served as the reference (Figure 3B). The treatments were categorized into three groups: the first group induced low pro-metastatic activity, lower than that induced by TCM. The second group displayed intermediate pro-metastatic activity, while the third group induced strong pro-metastatic activity, significantly exceeding that of TCM (Figure 3B). The most prevalent single treatments associated with a high transmigration score are ANGP2, VEGFC, and D while those associated with a low transmigration score are PLGF, EPO, and TGFB1 (Figure 3C). Of note, the prevalence of single treatments in low and high pro-metastatic activity groups (Figure 3C) did not reflect their transmigration OPLS weights as individual treatments (Figure 3A, triangle), suggesting intricate regulatory interactions between these treatments. Consistent with this, the inferred network revealed that these treatments were highly inter-connected with complex direct and indirect positive and negative regulations (Figure 2) highlighting the role of multiple interrelated pathways controlling TAM pro-metastatic activity.

**Figure 2.**
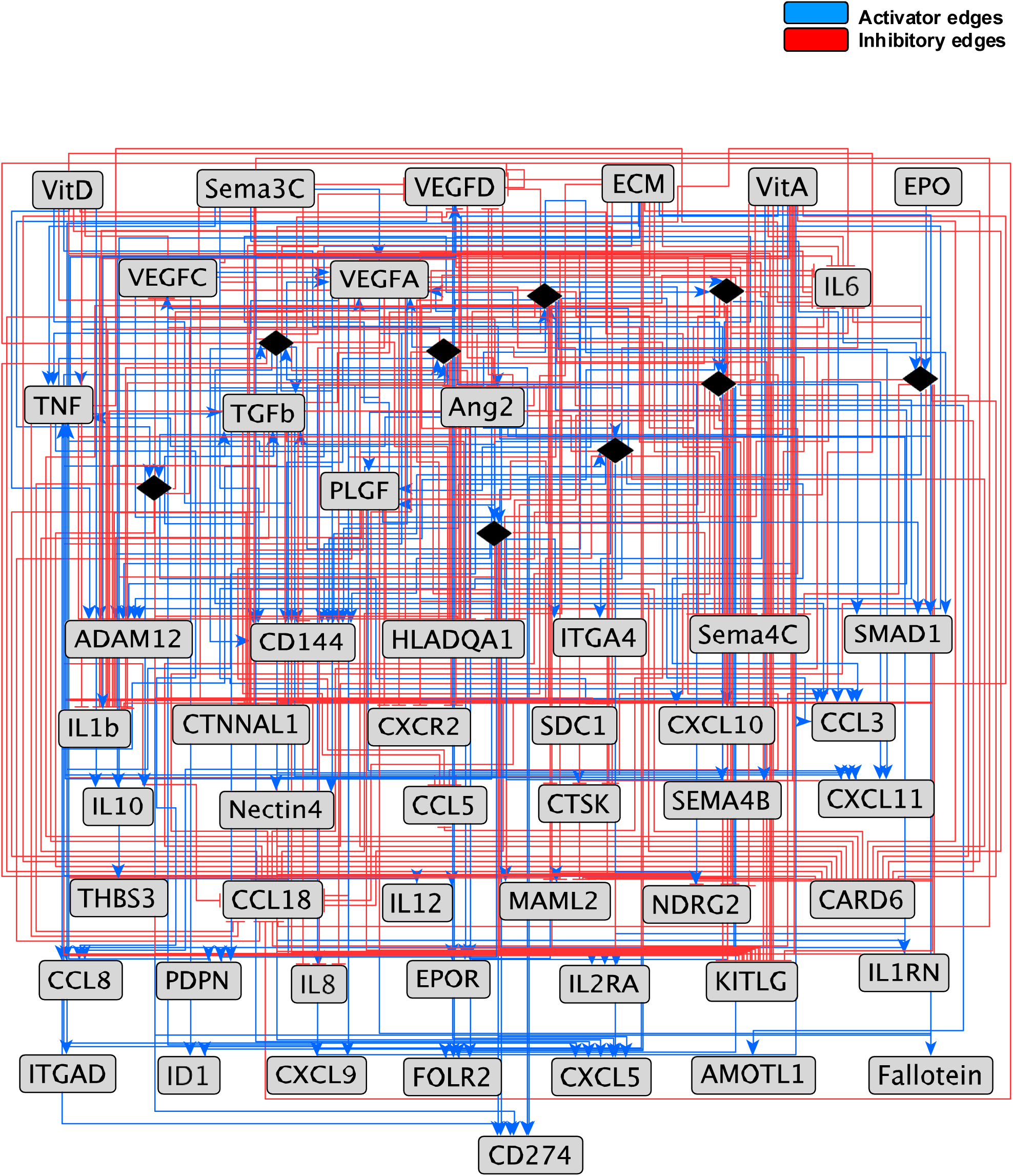
Inferred regulatory network of pro-metastatic TAMs based on experimental data. Inferred regulatory network of pro-metastatic TAMs based on experimental data. Node and edges activatory/inhibitory are shown in red and blue, respectively.

**Figure 3.**
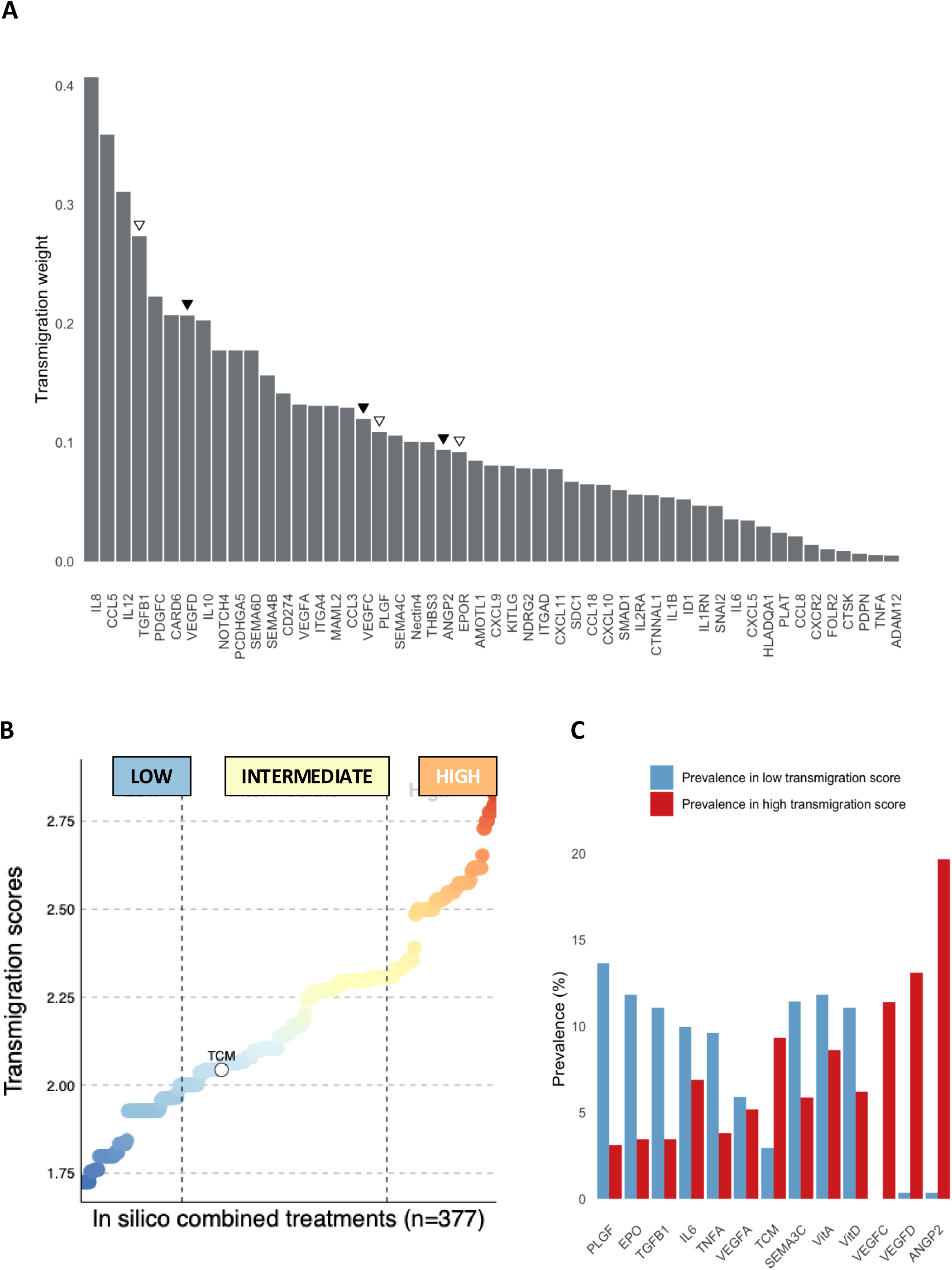
In silico predictions of the tumor cell transmigration activity. Calculation of the predicted transmigration score defined as the sum of the OPLS weights of active nodes of the network in response (A) to single, and (B) all single, double and triple treatments. Predicted transmigration scores ranked as low (lowest quartile), high (highest quartile) and intermediate transmigration. TCM induced an intermediate transmigration score and was used as a reference. (C) Prevalence of tested stimuli in combinations of treatments that induced in silico a low and high transmigration score.

**Table 1.** Experimentally tested perturbations and multi-omics features retained for network construction. In addition to performing a tumor cell transmigration assay with TAMs exposed to the treatments listed in the first column, the expression of the pro-metastatic marker was measured by FACS at the surface of TAMs (black, third column), a panel of soluble factors was quantified from TAMs culture conditioned medium (red, third column) and a panel of genes expression profiled in TAMs by real-time quantitative PCR (blue, third column).

| Perturbations and (concentration used) | Experimentally Tested Perturbations | Retained Perturbations | Retained functional features |
| --- | --- | --- | --- |
|  | Negative Control (plaint medium) | Negative Control (plaint medium) | CD144 |
|  | Positive Control TCM | Positive Control TCM | CD274 |
| Vascular Endothelium Growth Factor-A (2 ng/ml) | VEGF-A | VEGF-A | IL1RN |
| Vascular Endothelium Growth Factor-C (2 ng/ml) | VEGF-C | VEGF-C | Fallotein |
| Vascular Endothelium Growth Factor-D (2 ng/ml) | VEGF-D | VEGF-D | CCL8 |
| Vitamin A (10 ng/ml) | VitA | VitA | ADAM12 |
|  | VEGF-A+VEGF-C | VEGF-A+VEGF-C | CARD6 |
|  | VEGF-A+VEGF-D | VEGF-A+VEGF-D | AMOTL1 |
|  | VEGF-C+VEGF-D |  | TGF-β |
|  | VEGF-A+VitA |  | CTSK |
|  | VEGF-C+VitA |  | CXCL5 |
|  | VEGF-D+VitA | VEGF-D+VitA | FOLR2 |
| Vitamin D (1 μM) | VitD | VitD | HLADQA1 |
|  | VEGF-A+ VitD |  | ITGAD |
|  | VEGF-C+VitD | VEGF-C+VitD | CTNNAL1 |
|  | VEGF-D+VitD |  | SDC1 |
|  | VitA + VitD |  | ITGA4 |
| Colony Stimulating Factor-1 (20 ng/ml) | CSF-1 |  | ID1 |
| Interleukin-34 (20 ng/ml) | IL-34 |  | THBS3 |
| Granulocyte-macrophage colony-stimulating factor (5 μg/ml) | GM-CSF |  | PDPN |
| Tumor Growth Factor beta (100 ng/ml) | TGF-β | TGF-β | Sema4C |
|  | CSF-1+IL-34 |  | Nectin4 |
|  | CSF-1+GM-CSF |  | CXCR2 |
|  | CSF-1+TGF-β |  | SMAD1 |
|  | GM-CSF+IL-34 |  | IL2RA |
|  | IL-34+TGF-β |  | SEMA4B |
|  | GM-CSF+TGF-β |  | CCL18 |
| Angiopoietin -2 (100 ng/ml) | ANG-2 | ANG-2 | NDRG2 |
| Placenta Growth Factor-1 (10 μg/ ml) | PlGF-1 | PlGF-1 | KITLG |
| Tumor Necrosis Facctor-alpha (10 μg/ ml) | TNF-α | TNF-α | MAML2 |
| Prostaglandin E2 (100 ng/ml) | PGE2 |  | EPOR |
|  | ANG-2+PlGF1 | ANG-2+PlGF1 | VEGF-A |
|  | ANG2+TNF-α | ANG2+TNF-α | CXCL9 |
|  | ANG2+PGE2 |  | CCL5 |
|  | PlGF-1+TNF-α |  | CXCL11 |
|  | PlGF-1+PGE2 |  | CCL3 |
|  | TNF-α+PGE2 |  | CXCL10 |
| Interleukin-1 (10 μg/ ml) | IL1 |  | IL-6 |
| Interleukin-6 (10 μg/ ml) | IL6 | IL6 | IL-8 |
| Interleukin-4 (10 μg/ ml) | IL4 |  | IL-10 |
| Erythropoietin (10 U/ml) | EPO | EPO | IL-12 |
|  | IL1+IL6 |  | MCP-1 |
|  | IL1+IL4 |  | MMP9 |
|  | IL1+EPO |  | IL-1b |
|  | IL6+IL4 |  | IL-23 |
|  | IL6+EPO | IL6+EPO | TNF-α |
|  | IL4+EPO |  |  |
| Semaphorin 3C (200 ng/ml) | SEMA3C | SEMA3C |  |
| Semaphoin 3F (200 ng/ml) | SEMA3F |  |  |
|  | SEMA3C+SEMA3F |  |  |
|  | ANG2+TNF-α+PLGF-1 | ANG2+TNF-α+PLGF-1 |  |
|  | ANG2+PLGF-1 | ANG2+PLGF-1 |  |
|  | TNF-α+PLGF-1 | TNF-α+PLGF-1 |  |

### The inferred dynamical network of pro-metastatic TAMs is complex

Our inferred network is composed of 52 nodes, including 9 ‘AND’ gates and 381 directed and signed edges (inhibitory or activatory). Six treatment nodes -Vitamin D (VitD), Vitamin A (VitA), Erythropoietin (EPO), Tumor conditioned medium (TCM), Sema3C, and IL-6-act solely as inputs (in-degree = 0) within the network. Additionally, seven soluble factors (ANGP2, PLGF, TNFA, TGFB1, VEGFA, VEGFC, and VEGFD) function as intermediate nodes, meaning they are both measured as readouts and applied as treatments, enabling the capture of feedback loops essential for simulating dynamic TAM responses. The rest of the nodes are measured readouts and have an out-degree of 0. The inferred dynamical network (Figure 2) reveals a dense and highly interconnected structure, composed of multi-layered feedback loops and signaling cascades that regulate TAM function. Key network metrics (Supplementary Figure 1) further illustrate this complexity. Nodes such as TNF, ANGP2, VEGFA, and VEGFC exhibit high degrees, underscoring their central roles within the network and suggesting that they may act as major hubs in pro-metastatic signaling pathways. Additionally, betweenness centrality highlights ANGP2, VEGFC, and VEGFD as critical connectors within the network, directing the flow of information across various pathways. This level of complexity underscores the need for a computational framework capable of disentangling intricate pathways to identify relevant intervention points.

### Identification of critical pathway cross-talks underlying TAM pro-metastatic activity

We delineated the key biological pathways underlying TAM pro-metastatic activity by computing the transmigration scores of single, double, and triple treatment combinations. None of the single treatments alone scored higher than TCM and the breadth of single treatment scores was lower than the one of double treatments, which, in turn, was lower than the one of triple treatments (Figure 4A).

**Figure 4.**
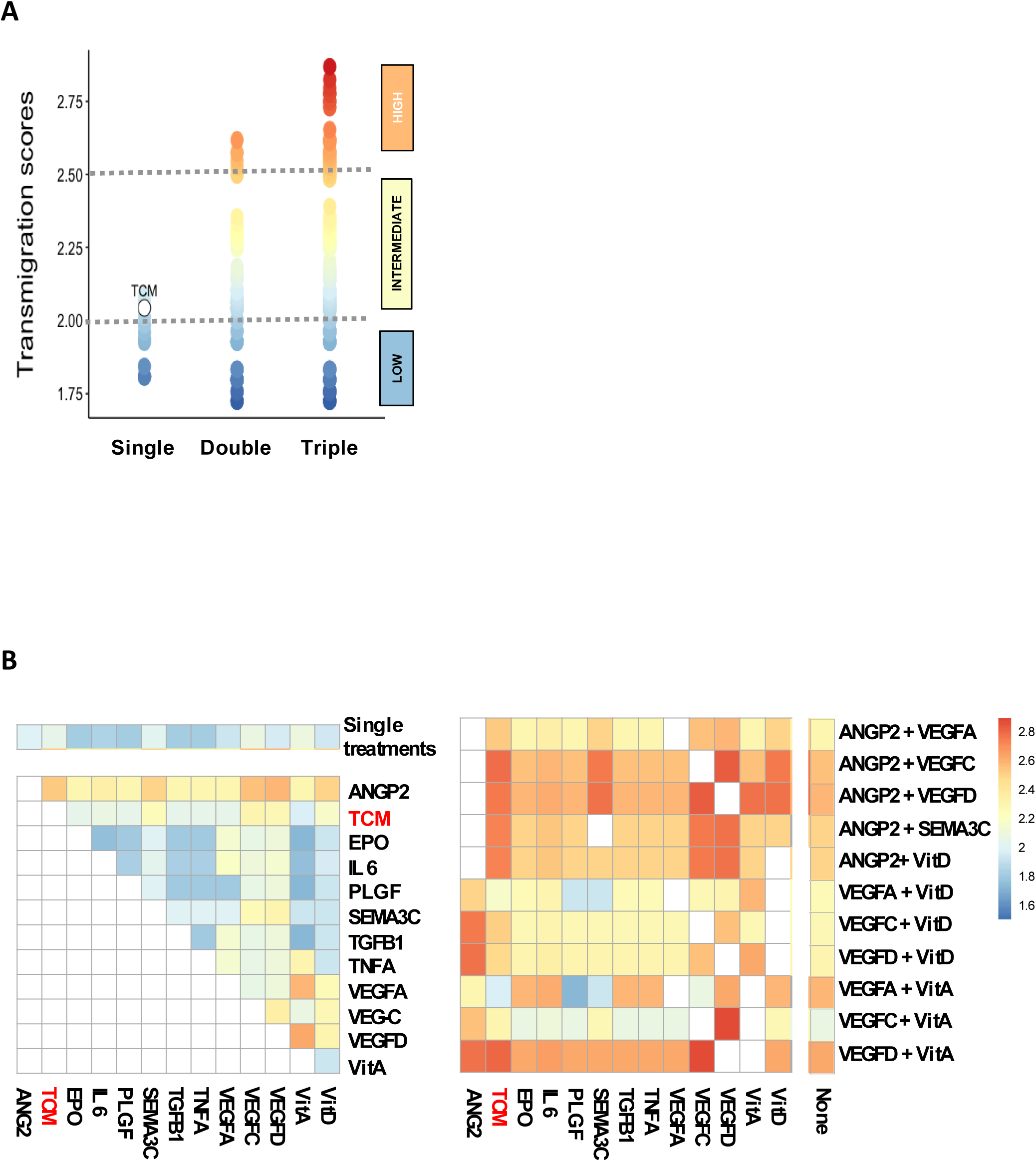
Identification of key pathways and pharmacological interventions underlying TAM pro-metastatic activity based on TAM dynamical network. (A) In silico predicted transmigration scores of single, double, and triple treatments. (B) Corresponding in silico predicted scores in a set of single, double, and triple treatments.

Importantly, when Ang-2 was combined with other treatments, it consistently enhanced TAM pro-metastatic activity which was always higher than TCM (Figure 4B, left panel). Notably, combinations of ANGP2 with VEGFC, VEGFD, SEMA3C, or VitD resulted in significantly higher scores, suggesting a strong synergistic interaction between Ang-2 and these axes (Figure 5B, left panel). A comparable synergy was observed between VitA and VEGFA/VEGFD axes (Figure 5B, left panel). Triple combinations of ANGP2, VEGFC, VEGFD, SEMA3C, and VitD further enhanced TAM pro-metastatic activity relative to double combinations except for VitA when combined with VEGFD/ANGP2 or SEMA3C/ANGP2. SEMA3C strongly increased the trans-migration score only when combined with ANGP2 and VEGFC or VEGFD (Figure 5B, right panel). Together these findings indicate that TAM pro-metastatic activity is governed by intricate cross-talks between TIE2 (receptor of ANGP2), VEGFR-2 and VEGFR-3 (receptors of VEGF-C and -D) pathways, and VitD/VitA pathways. In contrast, the VEGFR-1 pathway seems less critical, as ANGP2 when combined with VEGFR-1 ligands (i.e. VEGF-A and PlGF) resulted in a much weaker transmigration score.

**Figure 5.**
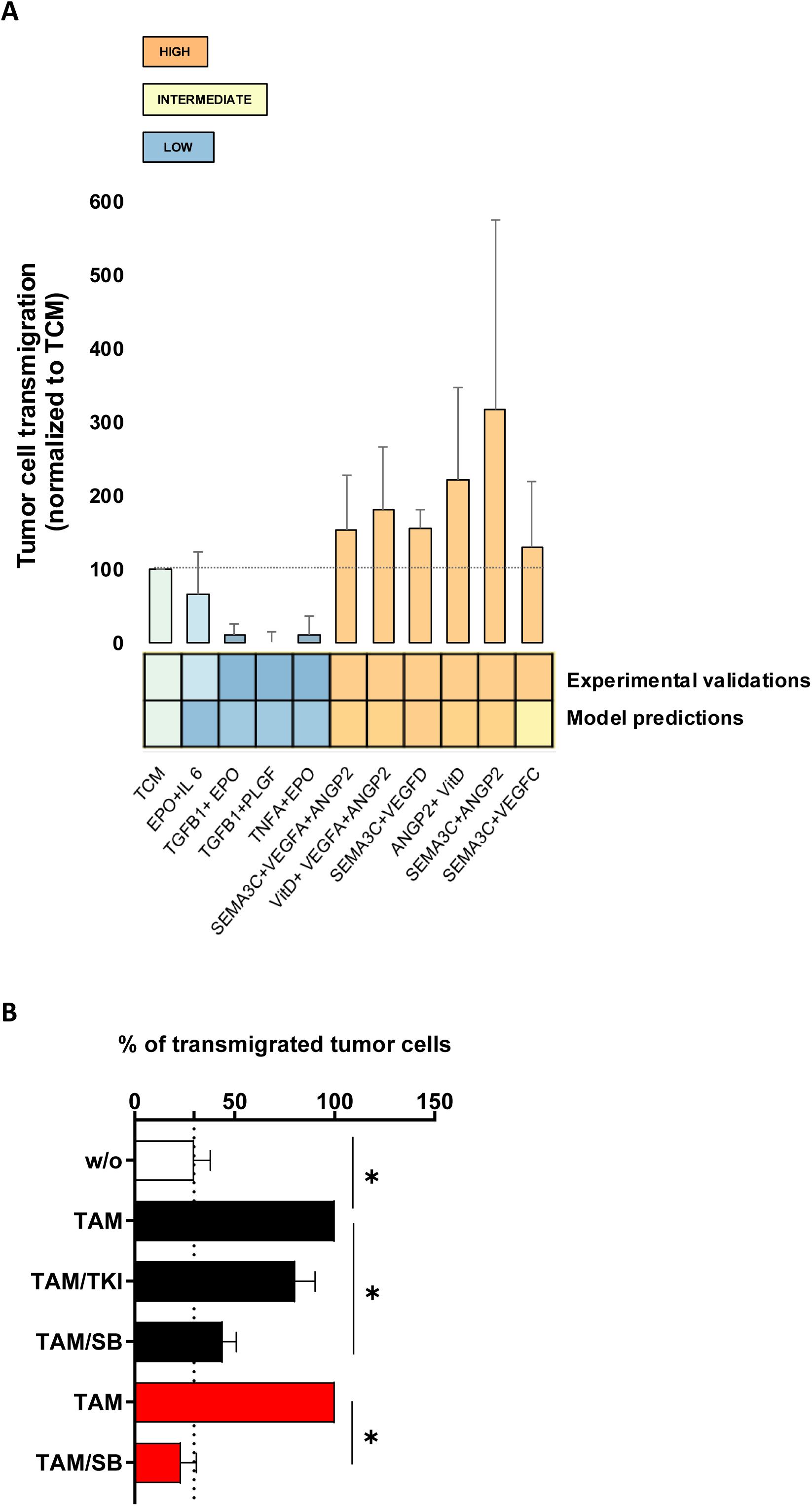
The regulatory network of pro-metastatic TAMs successfully simulates TAM behavior and identified powerful pharmacological interventions capable of dampening their pro-metastatic activity. (A) Experimental assessment in vitro of in silico predicted combined treatments associated with low and high transmigration scores of tumor cell transmigration, HV: High Variability. (B) Impact of pharmacological intervention on TAM pro-metastatic activity measured experimentally using the tumor cell transmigration assay. The assay was performed with TAM differentiated in vitro (black bars) and TAM isolated from patient tumors (red bars). TAM were either left untreated or treated with TKI and SB. A minimum of five distinct experiments were cumulated and the groups were statistically compared with Kruskal-Wallis test followed by Dunn’s multiple comparison post-test (*p<0.05).

### TAMs’ dynamical network reliably predicts TAM pro-metastatic activity

To evaluate the robustness and reliability of our model, we experimentally tested a set of ten combined treatments that were predicted in silico as spanning a broad range of transmigration scores (Figure 5A). For all these 10 treatments, the amplitude of the transmigration score predicted in silico was confirmed experimentally. However, two combinations (SEMA3C + ANGP2 and SEMA3C + VEGFC) exhibited a high experimental variability reflected by high standard deviations (Figure 5A). Of note, for each of these experiments, distinct individual blood donors were used from which monocytes were derived. This observation was reproduced when increasing the number of experiments suggesting that the system stabilized in multiple attractor states instead of a unique one, depending on donors. Hence, it is likely that our model did not capture the whole complexity and heterogeneity of the wiring of the underlying pathways. In summary, our results underscore the complexity of the TAM pro-metastatic regulatory network (Figure 2), yet confirm that, in most cases, the model accurately and reliably predicts pro-metastatic activity (Figure 5A).

### TAMs’ dynamical network successfully predicts pharmacological interventions that dampen **patient TAM pro-metastatic activity**

Based on our understanding of the pathway interplays controlling TAM pro-metastatic activity (Figure 4), we speculated that pharmacological interventions targeting TIE2, VEGFR-2, and VEGFR-3 pathways simultaneously -but not TIE2 and VEGFR-1 pathways-would effectively impair TAM pro-metastatic activity. We computed the transmigration scores of the corresponding interventions by assimilating inhibitions of the kinase activity of these receptors to an inactive state of the corresponding ligands. The transmigration score of active TIE2, VEGFR-1, 2, and 3 pathways (i.e. when ANGP2, PLGF, VEGFA, C, D, and SEMA3C are active in the network) was 3.02 while the score of with all these nodes off was 1.01. The computed transmigration score of TIE2 and VEGFR-1 inhibition (i.e. ANGP2, PLGF, and VEGFA off) was 1.80. Consistent with these predictions, we observed that TAMs treated with TIE2 and VEGFR-1-specific kinase inhibitors (TKI) failed to impair TAM pro-metastatic activity while SB, a kinase inhibitor targeting TIE2, VEGFR-2, and VEGFR-3, did effectively (Figure 5B, black and white bars). Most importantly, we next evaluated the impact of SB on patient TAM in situ. To this end, we isolated TAM from patient breast tumors, exposed them to SB, and measured their capability to induce tumor cell transmigration using the same experimental settings. We observed a very strong inhibition of patient tumor TAM pro-metastatic activity following SB treatment (Figure 5B, red and white bars). These results reinforce the critical role of the TIE/VEGFR-2/VEGFR-3 axes in controlling TAM pro-metastatic activity and extend the highly predictive value of our model to pro-metastatic TAM infiltrating patient breast tumors.

## DISCUSSION

This study demonstrates the power of integrating experimental data with computational modeling to predict and modulate the pro-metastatic activity of TAMs in human breast cancer. The dynamical Boolean model allowed for the exploration of multiple pathway interactions and provided valuable insights into TAM-mediated metastasis. Importantly, the combination of in silico predictions with experimental validations enabled the identification of a potent pharmacological intervention, which would have been impossible to uncover through traditional experimental approaches alone.

Building on our prior experience with robust dynamical models ^7,8^, this study incorporated several key modifications and integrated not only cell-associated and soluble proteins but also a select set of transcripts to better capture TAM behavior within the tumor microenvironment. This integration of proteomic and transcriptomic datasets, which operate at distinct scales and kinetics, added complexity and required additional experimental repetitions to ensure accuracy. Additionally, we used OPLS analysis to quantify the impact of treatments on tumor cell transmigration. This analysis helped identify the key functional features driving pro-metastatic activity, providing a quantitative measure that enhanced the overall predictive power of our model.

Our study highlights the potential for novel therapeutic strategies by co-targeting key pathways that regulate TAM pro-metastatic activity, such as TIE2, VEGFR-2, and VEGFR-3. While tumor heterogeneity presents a significant challenge for discovering effective therapeutic combinations, our computational modeling approach has provided insights into how these pathways interact dynamically. Instead of experimentally testing a vast number of combinations -294203 single, double, and triple combinations would have been required-, our model successfully identified a potent pharmacological intervention with only 19 single and 34 combined treatments, offering a streamlined approach to designing combination therapies and understanding the mechanism of action. An additional challenge in our approach resides in modeling a specific TAM population from patient tumors by integrating key pathways of this ecosystem without modeling the entire tumor microenvironment.

Our findings highlight the potential of systems biology approaches in oncology, particularly for the rational design of combination therapies. By identifying critical pathway cross-talks, this method can guide the selection of drug combinations that effectively target the metastatic processes. The use of such dynamical networks is expected to accelerate the development of precision oncology strategies, reducing the need for extensive experimental testing and focusing resources on the most promising therapeutic interventions.

While our model successfully predicted TAM pro-metastatic behavior and pharmacological interventions, there are inherent limitations due to the selection of perturbations based on prior knowledge. Although most in silico predictions were confirmed experimentally, two perturbations displayed high variability, likely due to the presence of multiple attractor states that the model was unable to fully capture. This variability may stem from the inherent heterogeneity of TAM populations and the current model’s limitations in reflecting the diverse responses of these cells to all perturbations. Expanding the model with additional nodes is expected to better represent TAM behavior and improve its predictive power. Future work should aim to extend the model to incorporate more comprehensive datasets, including full transcriptomic and proteomic profiles along with oncogenic genomic alterations. Additionally, expanding the model to include other components of the tumor microenvironment could provide a more holistic understanding of tumor progression and drug resistance mechanisms. Our long-term goal is to apply this approach to patient-specific models, allowing for the stratification of patients beyond a limited set of markers. These individualized models, contextualized to specific tumor environments, will be iteratively refined through experimental validation. Ultimately, this strategy will guide the development of personalized therapeutic interventions, tailored to the unique biology of each patient’s tumor, representing a step towards precision oncology.

## METHODS

### Experimental workflow

The workflow of combination of experimental and computational modeling approaches are shown in Figure 1. First TAMs differentiated in vitro were exposed to single and combined perturbations to gather sufficient experimental data at multiple levels (functional, phenotypical, and transcriptional) to identify the pathways underlying TAM pro-metastatic activity. The experimental data obtained on bulk monocyte populations (mixed pro-metastatic and non-pro-metastatic TAMs populations) were deconvoluted to construct a computation dynamical modeling of pro-metastatic monocytes. Simulations of the outcome of double and triple treatments on TAM pro-metastatic activity were conducted and a select set of them was retained for experimental assessment. Once validated, the final model was used to predict pharmacological interventions capable of dampening (conversely enhancing) TAMs pro-angiogenic activity (Figure 1). The selected perturbations, genes, and cell-associated and soluble proteins selected for multi-omics profiling were based on our limited knowledge of human breast TAMs and extended to data reported in the literature on mouse and human TAMs^6,13–29^.

### In-vitro differentiation of monocytes into pro-metastatic TAMs

We took advantage of the comparable transcription profiles and pro-metastatic activity of breast cancer patient TAMs and in-vitro differentiated TAMs that we had established (Pabois A., Crespo I. et al., manuscript in preparation). When not specified, in-vitro differentiated TAMs were used and obtained following exposure of CD14 monocytes immunomagnetically selected from healthy donor peripheral blood mononuclear cells (StemCell Technologies Inc., Vancouver, Canada) exposed for 2.5 days to MCF-7 and MDA-MB-231 tumor cell line conditioned medium (10% v/v each, hereafter called conditioned medium). The fraction of pro-metastatic monocytes ranged from 4 to 20% based on the expression of the pro-metastatic marker and their pro-metastatic activity assessed in vitro (next section) and in vivo (Pabois A., Crespo I. et al., manuscript in preparation).

### Experimental design: from perturbations to TAMs multi-omics profiling

We perturbed TAMs with 19 distinct treatments either alone or in 34 pairwise combinations plus one triple combination. These treatments include lymphangiogenic and immune suppressive factors in addition to pro-inflammatory and pro-tumoral factors (Table 1). The impact of these perturbations was measured at the phenotypical, functional (tumor cell transmigration assay and alteration of paracrine secretion profile of monocytes), and transcriptomic levels (we profiled changes in the expression of a select set of 40 genes differentially expressed in patient pro-metastatic TAMs. Briefly, monocytes were isolated by CD14 immunomagnetic positive selection (StemCell Technologies Inc.), exposed to treatments for 2.5 days and stained to analyze the frequency of pro-metastatic monocytes by FACS (based on the fraction of monocytes expressing the pro-metastatic marker), collected in RLT buffer for RNA extraction (Qiagen) and quantitative RT-PCR (10 000 cells), and harvested for transmigration assay (10 000 monocytes).

#### • Profiling of gene expression

RNAs were quantified by Quant-iT RiboGreen (Thermo Fisher Scientific), and their quality was assessed with a Fragment Analyzer (AATI), which showed an absence of RNA degradation. Reverse transcription was then performed following Fluidigm instructions (protocol 100-6472 B1), with a starting amount of 15 pg. A pre-amplification of 17 cycles was subsequently performed following Fluidigm instructions (protocol 100-5876 C2) with Fluidigm reagents and TaqMan assays (Thermo Fisher Scientific). The quantitative PCR was then performed with four 48.48 BioMark chips on a BioMark HD instrument following Fluidigm protocol for TaqMan assays (protocol 68000089 H1). Post-run quality control analysis and Ct calculation were performed automatically by the BioMark HD software, and were then confirmed manually. In parallel, monocyte conditioned medium was collected to profile paracrine secretions.

#### • Tumor trans-migration assay to measure TAM pro-metastatic activity

The aptitude of TAMs to induce the transmigration of tumor cells through the lymphatic endothelium was used as a metric of TAM pro-metastatic activity as this function correlated with the ability of in vitro differentiated TAMs and marker positive TAMs to induce tumor cell metastasis to the lungs in experimental mouse models of cancer (Pabois A., Crespo I. et al., manuscript in preparation). In tumors, the lymphatic endothelium that composes the lymphatic endothelium is leaky. We mimicked this condition by exposing a confluent layer of primary Human Dermal Lymphatic Endothelial Cells (HDLEC) grown at the surface of a transwell membrane and exposed to breast tumor cell line conditioned medium^30^). Last, GFP-positive breast tumor cells (GFP-recombinant MDAM-MB-231 breast tumor cells) are added to the upper chamber (Figure 1). In this assay, monocytes inserted into the lymphatic monolayer induce the transmigration of GFP-tumor cells to the lower chamber, which is monitored quantitatively by fluorescence microscopy (Figure 1). The monocytes were previously stained with a red fluorescent cell linker and their insertion into the lymphatic layer was monitored quantitatively by fluorescent microscopy of the surface of the transwell membrane (Figure 1).

For transendothelial cell migration assay, 20’000 HDLEC were seeded on gelatin-coated 24-well transwell inserts (8 µm pores; Corning Inc., NY, USA) and activated with conditioned medium. CD14+ monocytes (10 000) stained with a red fluorescent cell linker, PKH26 (Sigma-Aldrich), were added to the HDLEC monolayer for 5 hours to allow insertion into the lymphatic endothelium. 25 000 MDA-MB-231-GFP cells, grown overnight in RPMI 1640 with 2% FCS, were added to the culture for 48 hours in an IncuCyte live-cell analysis system (Essen Bioscience, Inc, Ann Arbor, MI). The number of inserted monocytes and transmigrated MDA-MB-231-GFP cells were counted with IncuCyte ZOOM software (Essen Bioscience). When indicated, TAMs were treated with TIE2 inhibitor compound 7 from Santa Cruz Biotechnology (Dallas, TX) and VEGFR inhibitor Vatalanib from Selleckchem (Houston, TX) and at 5 μM^31^. 4-(5-(6-methoxynaphthalen-2- yl)-2-(4-(methylsulfinyl)phenyl)-1H-imidazol-4-yl)pyridine (Sigma-Aldrich), named thereafter as SB27, was used at 0.25 μM.

#### • Profiling of TAMs functional phenotype by flow cytometry

Following blocking of Fc receptors with antibodies (Miltenyi Biotec, Bergisch Gladbach, Germany), cells were immunolabeled with the following anti-human mAbs: anti-CD45-Pacific orange, anti-CD11b-Pe-Cy7, anti-CD14-PerCP-Cy5.5 (Biolegend), anti-APC, anti-pro-metastatic marker-APC, Endoglin-PE (Clone 103, Thermo Fischer Scientific) and anti-Nrp2-Alexa Fluor 700 (Clone #257103, R&D Systems, Minneapolis, Minnesota, USA). Dead cells were excluded from analysis with a DAPI staining (Thermo Fisher Scientific). Fluorescence was measured on 50,000 cells/sample using a FACS LSR II® (BD Biosciences) equipped with a 610/20 nm filter on the violet detector and analyzed using FlowJo® 10 software (Tree Star, Inc., Ashland, OR).

#### • Profiling of TAMs paracrine secretion

Secreted cytokines and pro-angiogenic factors (Table 1) were measured and quantified in cell conditioned medium using AimPlex Multiplex assays (AimPlex Biosciences Inc., Pomona, CA).

### Deconvolution of Signals from Mixed Populations of Pro-Metastatic and Non-Prometastatic TAMs for Model Construction

Cumulated functional features (i.e. paracrine secretions and gene expressions) from TAMs were measured experimentally across the entire pool of stimulated cells, which includes a mixed population of pro-metastatic and non-prometastatic TAMS. To construct a regulatory network specific to pro-metastatic monocytes, it was necessary to extract subpopulation-specific information from this mixed data. This was achieved through computational deconvolution methodologies, which are designed to extract cell type-specific information from heterogeneous samples similar to the one we have previously reported^32^. The relative proportions of each monocyte subtype within the mixed population were determined using FACS. Subsequently, partial deconvolution analysis was performed using the Lawson-Hanson algorithm for non-negative least squares (NNLS), as implemented in the R package “nnls.”This allowed for the partial deconvolution of experimental readouts, enabling the differentiation of signals contributed by pro-metastatic versus non-prometastatic TAMs.

### Model construction and simulation

We constructed a regulatory network in which nodes represent either treatments or functional features, and edges denote the activation or inhibition of these functional features in response to perturbations. In some cases, perturbation targets were also measured as functional features, enabling the incorporation of feedback processes into the model. To establish edges between treatments and functional features, we employed three key criteria: amplitude, reproducibility across replicates, and coherence across experimental conditions. A treatment was considered significant if it induced a change of sufficient amplitude, specifically when the perturbation effects on functional features were within the upper (indicating an increase in gene or soluble/cellular factor expression) or lower (indicating a decrease) quartiles of variation.

The control treatment, consisting of plain RPMI 1640 medium containing 10% FCS, served as the reference unperturbed state, against which the impacts of all other perturbations were evaluated. A link was considered reproducible if the perturbation induced consistent effects in at least two-thirds of the biological replicates. To categorize these effects in a Boolean framework, we classified them into three categories: 1) no impact, where no link was established between the treatment and the functional features; 2) an increase in the readout relative to the unperturbed state, leading to the addition of an activatory edge (“->”); and 3) a decrease, indicating inhibition relative to the unperturbed state, leading to the addition of an inhibitory edge (“-|”).

Furthermore, only coherent links were retained—those where the connected nodes exhibited consistent trends across treatments, whether correlated or anti-correlated. The resulting dynamic network is illustrated in Figure 2. To simulate the effects of treatments, we utilized BoolSim with an asynchronous updating scheme, allowing us to explore the impact of individual treatments as well as all possible combinations of double and triple treatments. We then analyzed the attractors—stable states of the network reached after perturbations—to assess their effects on the transmigration phenotype.

It is worth noting here that the added value of the modeling part in this work lies in its ability to capture the cross-regulation between genes downstream of two given perturbation nodes, which can modify the behavior of these nodes in combined treatments relative to single treatments. By limiting ourselves to the Boolean stable states of the system under perturbation, we can capture this cross-regulation. This allows for the creation of rankings and prioritization of specific combinations, providing a useful tool for identifying relevant intervention points in the complex pro-metastatic signaling pathways of TAMs.

Several limitations should be noted regarding our model. First, the predictive capabilities are confined to the perturbation space of the selected single treatments, restricting the model’s scope to these perturbation nodes. Secondly, feedback loops are only possible in nodes that serve as both perturbation and readout nodes, limiting the model’s ability to capture cross-regulation beyond these dual nodes. Additionally, the logic between coregulators is not inferred; an inhibitory dominant system is adopted as default logic, which may not accurately reflect the relationships between activators and inhibitors for all nodes, potentially altering the attractors. Finally, it is worth noting here that the assessment of combined perturbations is based on reachable stable states, some of which may be difficult to achieve and not significantly occupied.

### Orthogonal Partial Least Squares (OPLS) to calculate attractor transmigration scores

To score the capacity of simulated perturbations to induce monocyte pro-metastatic activity (i.e. the capacity of monocytes to induce transmigration of tumor cells across the lymphatic endothelium which was used as a metric of pro-metastatic activity), we first estimated the relative contribution of the functional features to transmigration by using Orthogonal Partial Least Squares (OPLS) multivariate analysis. The transmigration assay was used to quantify the effect of each treatment on the capability of monocytes to induce tumor cell transmigration across the lymphatic endothelium. Tumor Conditioned Medium (TCM)-treated monocytes were used for normalization (100%). OPLS allows removing the dispersion that is not associated with transmigration and assigns a relative higher contribution to functional features more strongly associated to transmigration. We extracted the OPLS weight for each functional feature, which represents its contribution to the correlated variation to transmigration. Then, we calculated, for each attractor of the network under a simulated perturbation, a score consisting of the sum of OPLS weights of active nodes (functional features). A low score indicates a low transmigration and *vice versa*. Since each perturbation could lead to multiple attractors, for each perturbation we chose the attractor with the maximal transmigration score (Figure 3B).

From the OPLS analysis, we extracted weights for each node in the network, representing its relative contribution to the observed variation in transmigration. For each attractor identified in the network under simulated perturbation, we calculated a transmigration score by summing the OPLS weights of the active nodes (genes/cytokines). The score is given by the formula: *Score* = 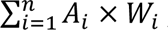, where *A*_*i*_ denotes the activation status of each gene/cytokine (1 for active, 0 for inactive) and *W*_*i*_ represents its OPLS-derived weight. A lower score corresponds to reduced transmigration activity, and vice versa. Since each perturbation may result in multiple attractors, we selected the attractor with the highest transmigration score for further analysis, as this approach showed better correlation with experimentally observed transmigration effects induced by double treatments (correlation coefficient: 0.756) compared to the minimal attractor transmigration scores (correlation coefficient: 0.5).

### Challenging experimentally model predictions to assess model reliability and accuracy

Predicted transmigration scores of all in silico single, double, and triple treatments were computed and ranked as low (lowest quartile), high (highest quartile), and intermediate transmigration. Tumor Conditioned Medium (TCM) induced an intermediate transmigration score which was used as a reference (Figure 3B and 4). A select set of combined treatments was tested experimentally to assess the validity and accuracy of the model (Figure 5A).

## ACKNOWLEDGMENTS

The authors thank Vérène Pignat for her excellent technical work. This work was supported by grants from the Swiss National Foundation: M.-A. Doucey, project 310030-120473.

---

**Supplementary Figure 1:**
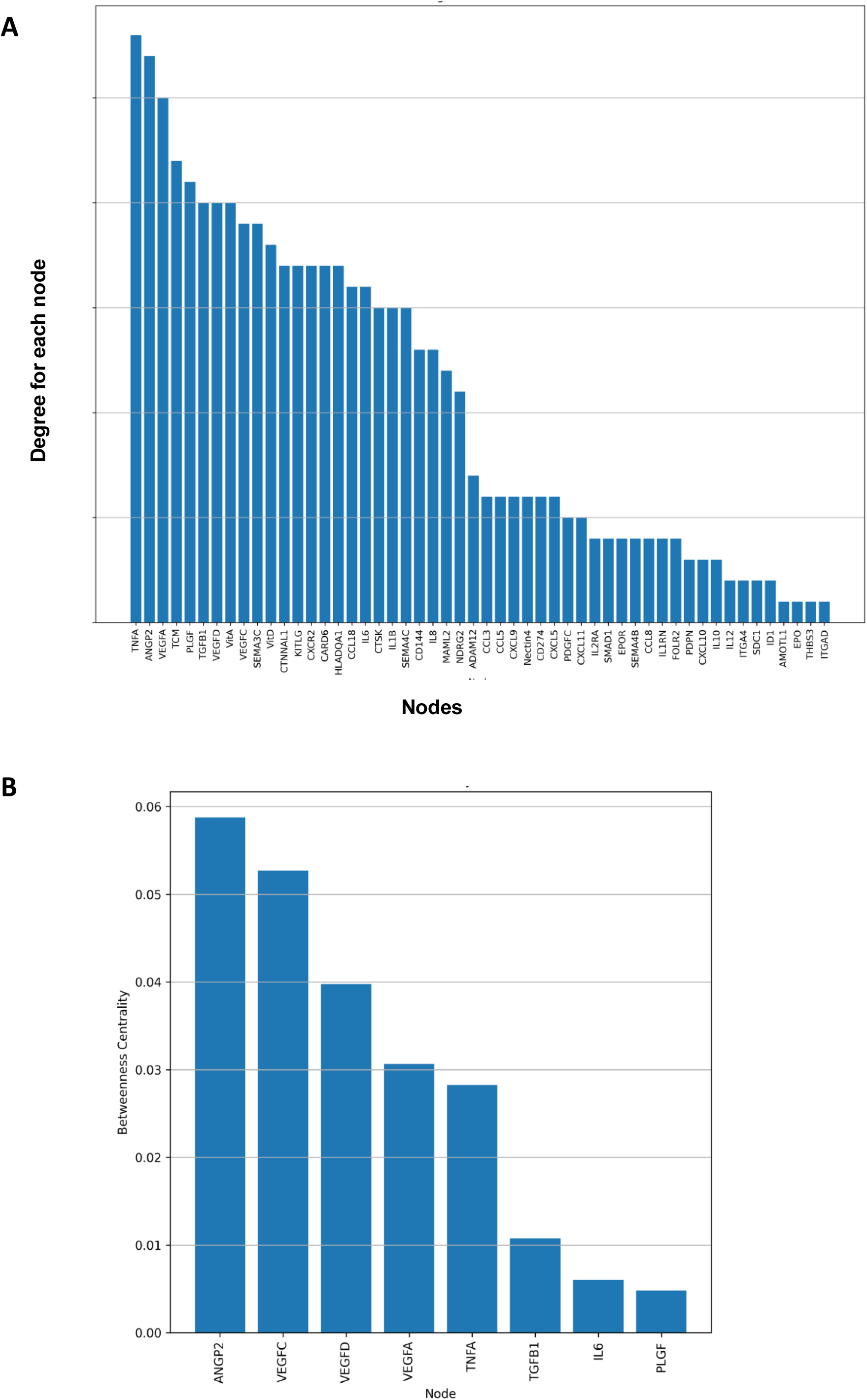
Degree and Betweenness Centrality of the TAM network nodes: (A) Bar chart of the degree of distribution of each node in the network, ordered from highest to lowest, indicating the number of connections each node has. (B) Betweenness centrality: Bar chart of the betweenness centrality values for each node, ordered from highest to lowest, highlighting nodes that frequently lie on the shortest paths between other nodes. Nodes with centrality 0 are not shown.

